# Assessment of brain-derived extracellular vesicle enrichment for blood biomarker analysis in age-related neurodegenerative diseases: An international overview

**DOI:** 10.1101/2023.10.02.560210

**Authors:** AmanPreet Badhwar, Yael Hirschberg, Natalia Valle Tamayo, M. Florencia Iulita, Chinedu T. Udeh-Momoh, Anna Matton, Rawan M. Tarawneh, Robert A. Rissman, Aurélie Ledreux, Charisse N. Winston, Arsalan S. Haqqani, Alzheimer’s Association International Society to Advance Alzheimer's Research and Treatment, BBB-EWG

## Abstract

**INTRODUCTION:** Brain-derived extracellular vesicles (BEVs) in blood allows for minimally-invasive investigations of CNS-specific markers of age-related neurodegenerative diseases (NDDs). Polymer-based EV- and immunoprecipitation (IP)-based BEV-enrichment protocols from blood have gained popularity. We systematically investigated protocol consistency across studies, and determined CNS-specificity of proteins associated with these protocols.

**METHODS:** NDD articles investigating BEVs in blood using polymer-based and/or IP-based BEV enrichment protocols were systematically identified, and protocols compared. Proteins used for BEV-enrichment and/or post-enrichment were assessed for CNS- and brain-cell-type-specificity; extracellular domains (ECD+); and presence in EV-databases.

**RESULTS:** 82.1% of studies used polymer-based (ExoQuick) EV-enrichment, and 92.3% used L1CAM for IP-based BEV-enrichment. Centrifugation times differed across studies. 26.8% of 82 proteins systematically identified were CNS-specific: 50% ECD+, 77.3% were listed in EV- databases.

**DISCUSSION:** We identified protocol steps requiring standardization, and recommend additional CNS-specific proteins that can be used for BEV-enrichment or as BEV-biomarkers.

## 1. INTRODUCTION

The number of individuals living with neurodegenerative conditions has more than doubled from 1990 to 2016, owing to population aging and growth [1,2]. By 2050, more than 150 million are expected to live with neurodegenerative diseases of aging (NDDs), with Alzheimer’s disease (AD) being the most prevalent [1]. NDDs have a long pre-symptomatic stage during which neuropathological changes occur prior to symptom onset [3,4]. While neuroimaging and cerebrospinal fluid (CSF) biomarkers have been established for AD diagnosis [5], they are expensive and/or invasive and available only in specialized centers. Major efforts are being devoted to the development of reliable early diagnostic biomarkers to facilitate identification of at-risk persons prior to symptom-onset. Availability of minimally-invasive blood biomarkers for early diagnosis and/or investigating disease mechanisms and potentially novel therapeutic targets would be of great value.

While blood collection is easy to perform, measurement of brain-derived markers from blood poses a challenge due to the complex composition of blood itself, and the relatively low quantities of molecules released from the brain into the peripheral circulation [6]. Technical and analytical advances over the past decade are, however, starting to enable specific and sensitive measurements of a handful of NDD-biomarkers in blood, with most being relevant to AD (e.g., Aβ, and specific tau phosphorylated forms) [7–11]. Despite this advancement, many of the current blood NDD biomarkers are not brain-specific, but instead also expressed at high levels in peripheral tissues (e.g. Aβ expression in red blood cells and cardiomyocytes) [12,13]. This renders the interpretation of blood-based measurements challenging. Approaches that allow for the enrichment of brain- and brain-cell-specific biomarkers from blood are thus being actively sought after.

On this front, extracellular vesicles (EVs) in blood comprise a promising minimally-invasive biomarker source for many diseases, including cancer and NDDs [14,15]. Released by all cells in the body [16], EVs are lipid-delimited nanoparticles of different intracellular origins, with cell-cell communication being a main function [17]. EVs contain molecules (e.g. proteins) that mirror the parental cell content and expression level, thereby providing a snapshot of the homeostatic status of their cell of origin. Given their small size, EVs can diffuse into biological fluids (e.g. blood, CSF (see review [18]), and bidirectionally cross the blood-brain barrier [19]. Their ability to diffuse from CNS to blood, from where they can be isolated, make them an attractive resource for novel biomarker discovery and mechanistic insights into brain disease.

Enrichment of both EVs and brain-derived EVs (BEVs) from blood is not without its challenges given the high non-EV content of blood (e.g. serum/plasma proteins, liposomes) [20]. Presently, several methods for enrichment of BEVs from blood exist, each with its advantages and disadvantages [15,21]. Of these, polymer-based precipitation followed by immunocapture with specific antibodies, commonly known as the immunoprecipitation (IP)-based BEV enrichment method, is widely used in NDD research [22]. While Fiandaca et al. [23] using the L1CAM antibody pioneered this method for enrichment of neuron-derived EVs (NEVs), others have since adapted/optimized it for enrichment of other brain-cell-type-derived EVs using other antibodies (e.g. glial) [22,24–28]. Given the growing use of IP-based BEVs enriched from blood in NDD research, the objectives for this study were to: 1) systematically assess protocols for IP-based BEV enrichment from blood, and 2) assess CNS-specificity and extracellular accessibility of proteins used for BEV enrichment and/or detected in BEV enriched isolates. Note that this study was conducted by the Alzheimer’s Association International Society to Advance Alzheimer’s Research and Treatment: Biofluid Based Biomarkers Professional Interest Area - Exosome Working Group (ISTAART-BBB-PIA-EWG).

## 2. METHODS

### 2.1 Literature Search

A comprehensive PubMed review on BEVs enriched from blood in AD had identified 26 articles published up to October 2019 [22]. We extended this search to March 22 2022, using six searches (Supplementary_Fig_S1). In addition to AD, we comprehensively reviewed the literature on BEVs enriched from blood in (a) other NDDs, namely, Parkinson’s disease, vascular dementia, frontotemporal dementia, and Lewy body dementia, and (b) Down syndrome - given that these individuals represent the largest population at genetic-risk for AD [29] (searches in Supplementary_Fig_S1). Given that the study of BEVs enriched from blood is an emerging field in NDD research, we conducted an additional 18 searches using less stringent keywords to avoid missing relevant studies (Supplementary_Fig_S1). In total, a combination of 54 keywords were used (Supplementary_Fig_S1).

Overall, only original research articles published in English and employing an IP-based BEV enrichment method from human blood were included. Additional articles were identified by scanning the reference lists of included articles. Articles excluded were: reviews/case-reports; those not investigating BEVs in NDDs. Those investigating BEVs in biofluids other than blood; non-human samples; and cell culture.

### 2.2 Data Extraction

Data extracted from articles that met inclusion/eligibility criteria were as follows: cohort demographics (number of participants, sex distribution); last-author demographics and publication date; blood-portion (plasma or serum) used for BEV-enrichment; protocol preparing plasma/serum for downstream EV and/or BEV work; E- enrichment protocol details; IP-protein used for BEV-enrichment and other protocol details; EV and BEV validation methods; and generating a list of proteins used for BEV-enrichment and/or investigated post-enrichment. For articles that provided partial or no protocol details, the published protocol(s) referenced by these articles was used to gather these details.

### 2.3 Characterization of Proteins in Our List

Proteins in our list (list generation described in Section 2.2) were converted to their corresponding UniProt [30] gene names to allow mapping to public databases. To determine CNS-specificity, “RNA consensus tissue gene data’’ from The Human Protein Atlas (v21.0 and Ensembl v103.38; proteinatlas.org, [31]) was utilized. It includes gene expression, corresponding to normalized expression values (nTPM), from 61 regions: 12 CNS and 49 peripheral regions (listed in Supplementary_Fig_S2). CNS specificity of each gene was calculated by comparing its expression levels in the 12 CNS regions with those in the 49 peripheral regions. Genes demonstrating ≥4-fold higher average in CNS than peripheral regions, and showing a statistically significant p-value <0.05 between the two groups (Mann-Whitney U-test) were defined as ‘CNS-specific’. Extracellular domain-containing (ECD) proteins, as potential targets for IP-based BEV-enrichment, were identified from the UniProt human database (v2022_01) [30]. EV specificity was identified using Exocarta (exocarta.org) and Vesiclepedia (microvesicles.org) databases. To determine brain-cell-type-specificity, a single-cell RNAseq dataset (accession GSE67835, [32]) reporting gene expression in six brain-cell-types in humans was downloaded from the NCBI GEO repository [33]. The six brain-cell-types were: neurons, oligodendrocyte progenitor cells (OPC), oligodendrocytes, astrocytes, microglia, and brain endothelial cells. For each identified CNS-specific gene, the cell-type-specific gene expression value was extracted from the GSE67835 dataset and normalized to the average expression among the six cell-types. If a gene had a normalized value ≥4-fold higher for a particular cell-type, it was tagged as specific to that cell-type, else it was tagged as present in multiple cell types.

## 3. RESULTS

### 3.1 Literature Search and Identification of IP-based BEV articles

Our search workflow identified 39 articles (53.4%) investigating BEVs enriched from blood using IP-based methods (Fig. 1A), of which 35 (89.7%) performed ELISA-based assays following enrichment. Most articles (64.1%) investigated AD or Parkinson disease (25.6%). Publication dates of articles ranged from 2014-2021, with most (28.2%) published in 2020 (Fig. 1B), and 55.1% reporting a last-author in the United States (Fig. 1C).

**Figure 1:**
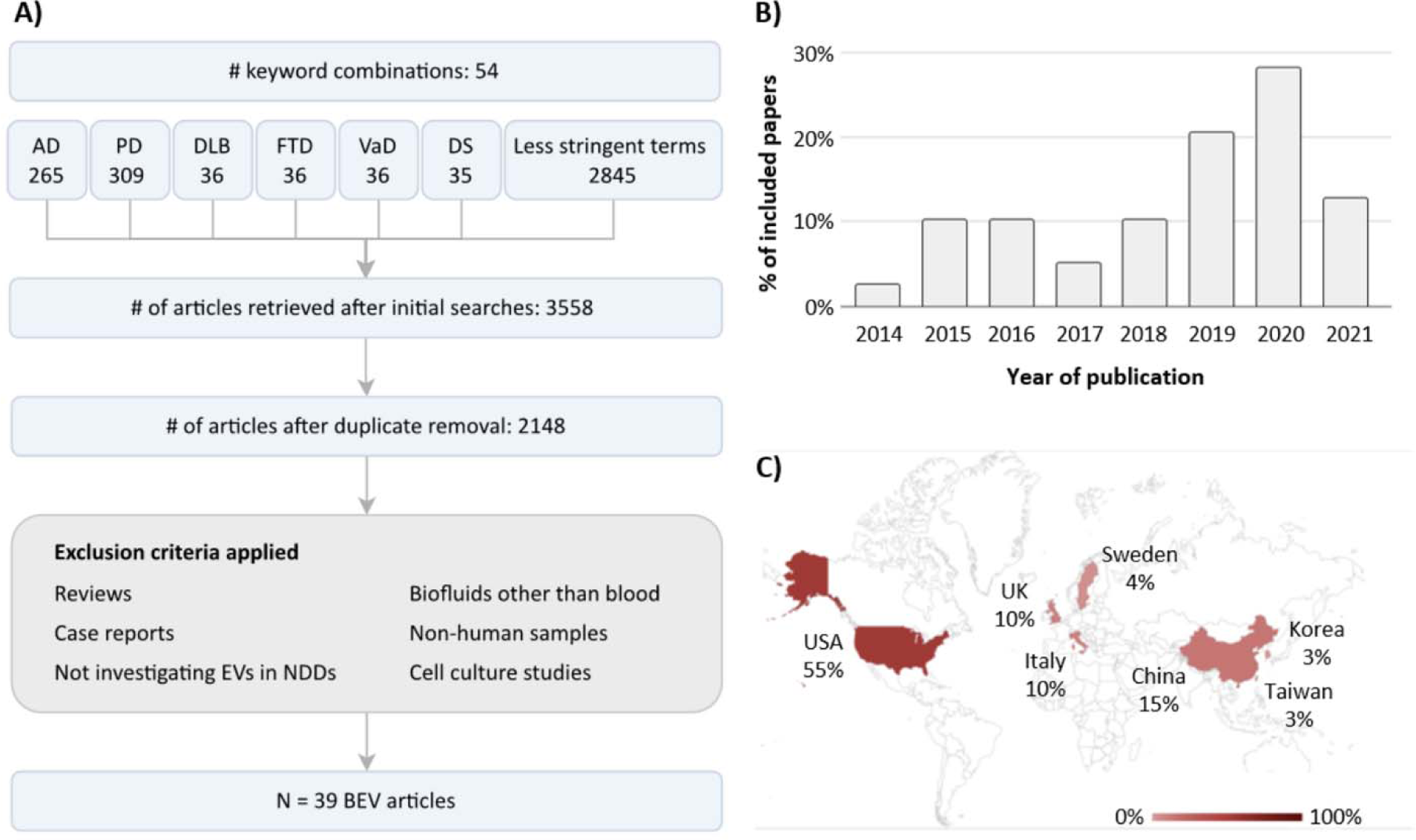
Literature Search (A) Flowchart of the article selection process. (B) Distribution of publication dates of the 39 BEV articles. (C) Geographic distribution of last-authors’ affiliations of the 39 IP-based BEV enrichment articles. Abbreviations: AD, Alzheimer’s disease; PD, Parkinson’s disease; DLB, Lewy body dementia; FTD, frontotemporal dementia; VaD, vascular dementia; DS, Down syndrome; EV, extracellular vesicles; BEV, brain-derived extracellular vesicles.

### 3.2 Assessment of Sample Preparation, EV and IP-based BEV Enrichment Protocols

Fig. 2 provides a graphical breakdown of protocols used for sample preparation, and EV and IP-based BEV enrichment.

**Figure 2:**
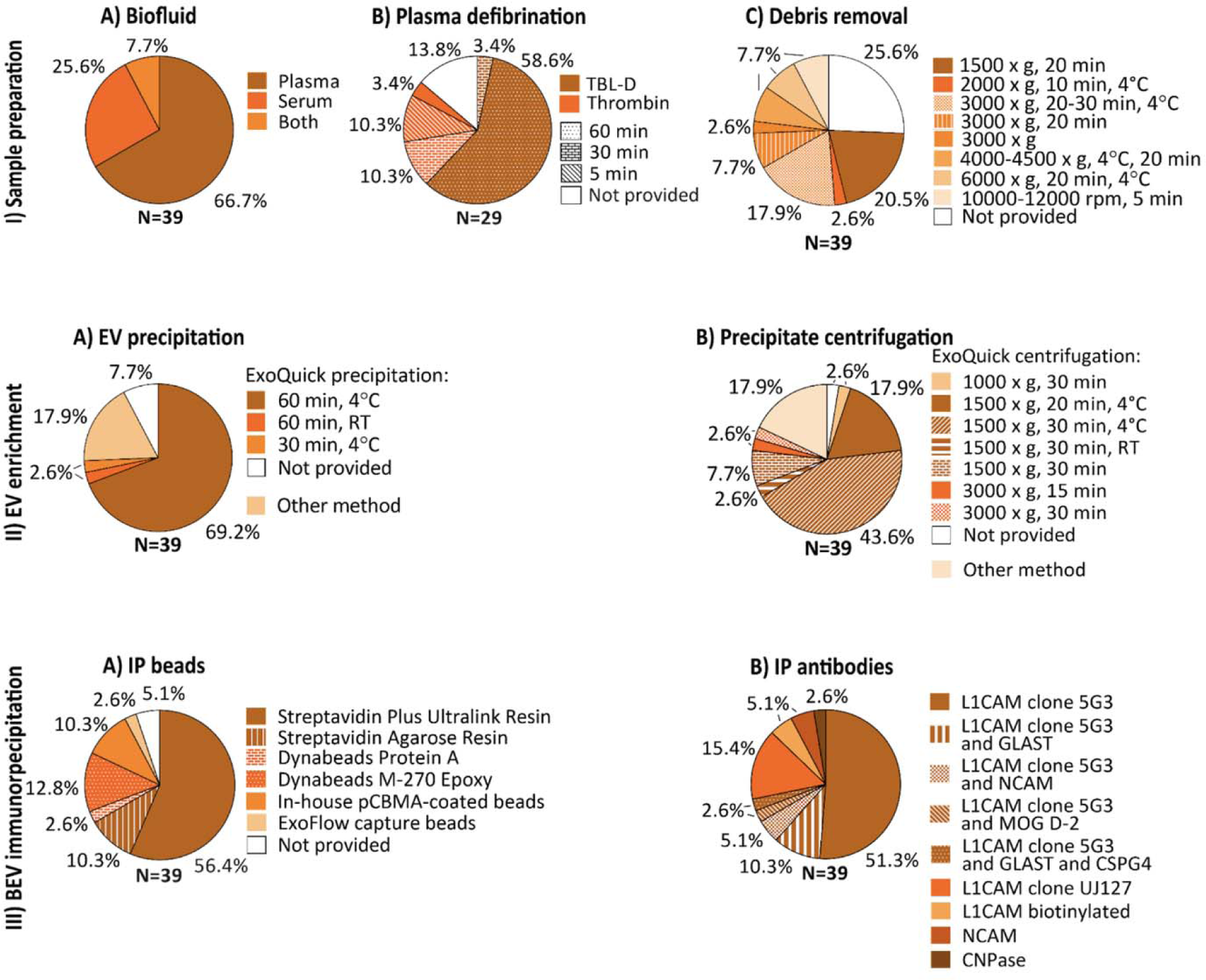
Sample preparation, EV and IP-based BEV enrichment protocols used. **(II) Sample preparation:** A) Biofluid used; B) Plasma defibrination - reagent used and incubation time; and C) Centrifugation for debris removal - speed, time and temperature. **(II) EV enrichment:** A) Method used for EV precipitation; for ExoQuick-based precipitation: incubation time and temperature; B) Centrifugation for EV precipitation - speed, time and temperature; **(III) BEV immunoprecipitation (IP):** A) IP beads used; B) Antibodies used for IP. Abbreviations: TBL-D, thromboplastin-D.

#### 3.2.1 Sample preparation for downstream enrichment of BEVs

The initial biofluid (plasma or serum) volume ranged from 250-500 μL across the 39 studies, with 26 (66.7%) using plasma only, 10 (25.6%) serum only, and three (7.7%) both (Fig. 2-IA). Of the 29 studies using plasma, 25 (86.2%) reported a defibrination-step using thromboplastin-D (N=18, 62.1%, 100-200 μL) or thrombin (N=7, 24.1%, 2.5-5 μL), while the remaining four did not mention this step (Fig. 2-IB). All studies using thromboplastin-D incubated for 60 mins at room temperature (RT), except for one study (30 mins, RT) (Fig. 2-IB). For thrombin, while an equal number of studies (N=3 each) incubated for 5 or 30 mins at RT (Fig. 2-IB), one study lacked specifics. Following incubation, all studies using thromboplastin-D, and four studies using thrombin added protease and/or phosphatase inhibitor cocktails diluted in Dulbecco’s phosphate buffered saline (DPBS, 150-495 μL). While a defibrination step is not needed for serum, five such studies also added DPBS (150-500 μL).

All 39 studies performed a centrifugation step to remove fibrinogen clot (relevant for plasma EV- studies) and/or cell debris (Fig. 2-IC). Centrifugation conditions (speed, time) varied between studies (Fig. 2-IC). A speed of 3000 x g, 20-30 min (N=12 studies, 11 plasma) was most used, followed by 1500 x g, 20 min (N=9 studies, 8 plasma). Centrifugation temperature was not reported by the majority of studies (N=24, 61.5%), but when reported (N=15, 38.5%), it was 4L (Fig. 2-IC).

#### 3.2.2 EV enrichment

Thirty-two (82.1%) studies performed polymer-based EV precipitation on defibrinated plasma or serum using ExoQuick^®^ (Fig. 2-IIA). Of these, 28 studies incubated with ExoQuick for 60 mins (temp: N=27, 4°C or on ice, N=1, RT), one for 30 mins (at 4°C), and three did not report incubation conditions (Fig. 2-IIA). For precipitation of EVs following incubation, most ExoQuick studies (N=28, 71.8%) centrifuged at 1500 x g for 20-30 mins, with 24/28 performing this step at 4°C (Fig. 2-IIB). Precipitated EVs were resuspended in solution containing protease/phosphatase inhibitor cocktails under continuous rotation. Of the seven (17.9%) studies not using ExoQuick, one used a combination of EV precipitation and size exclusion chromatography, three used sequential spins for EV enrichment, and three others did not perform EV enrichment.

#### 3.2.3 IP-based BEV-enrichment

All 39 studies performed IP-based BEV-enrichment. The majority of studies used anti-L1CAM products (36/39, 92.3%), albeit different types, to enrich NEVs (Fig. 2-IIIB). Of these, eight studies (22.2%) used an additional sample to enrich NEVs or other brain-cell-type-derived EVs with another antibody: NCAM (for NEVs), MOG (for oligodendrocyte-derived EVs), GLAST (for astrocyte-derived EVs), or CSPG4 (for CSPG4-cell-derived EVs). Resultant BEV preparations were incubated with magnetic or resin beads. Twenty-two (56.4%) of the studies used Streptavidin Plus UltraLink Resin beads (Fig. 2-IIIA) for immunoprecipitation.

### 3.3 EV and BEV Characterization by Size, Shape, and Cellular Origin

Twenty-four (61.5%) studies reported assessing biophysical properties of EVs and/or BEVs. Techniques most used were nanoparticle tracking analysis (NTA) and transmission electron microscopy (TEM), employed by 18 (46.2%) and 16 (41.0%) studies, respectively. Thirty-three (84.6%) studies reported assessing the presence of EV-specific markers, with tetraspanin protein marker CD81 (N=24, 61.5%) and the intracellular marker Alix (N=10, 25.6%) being most used. Other EV-markers used for validation were CD63 (N=6, 15.4%), CD9 (N=6, 15.4%), TSG101 (N=5, 12.8%) and Hsp90 (N=1, 2.6%).

Neuronal origin of BEVs were validated by 12 different markers: L1CAM, NCAM, NfL, neuronal specific (NS)-enolase, synaptophysin, Tau-1, MAP-2, NeuN, Enolase-2, MAPT, GRIA1, and PLP1. Of these, L1CAM (N=16, 41.0%), followed by NCAM (N=5, 12.8%) and NfL (N=4, 10.3%) were most used. Glutamine synthetase (GluSyn) was most used to validate astrocyte-derived EVs (AEVs). Studies using MOG or CNP-ase as an IP antibody, validated the presence of these proteins post IP (e.g., using ELISA). Fourteen studies (35.9%) did not mention any brain cell marker or provide a reference paper for its validation.

### 3.4 CNS-specificity, ECD status, and presence in public EV-databases of proteins in our list

We generated a list of 117 proteins that were used for BEV enrichment and/or investigated post-enrichment by the 39 articles, and these corresponded to 87 unique genes (Table 1). Protein products of five of these genes are known EV markers, namely, CD81, CD9, CD63, PDCD6IP (or Alix), and TSG101. CNS versus peripheral gene expression levels analysis on the remaining 82 genes identified 22 (26.8%) to be CNS-specific (Fig. 3A). Of these, 11 had ECDs (MOG, SYT1, SYP, SLC1A3, NRXN2, STX1A, NCAM1, L1CAM, NLGN1, GRIA4, SYT2), and 11 did not (GFAP, SNAP25, GAP43, NEFL, UCHL1, MAPT, ENO2, SNCA, OMG, NRGN, SYN1). Relative expression levels across the 12 CNS and 49 peripheral regions of the 17 CNS-specific genes whose protein products are known to be present in EVs are shown in Fig. 3B-D.

**Figure 3:**
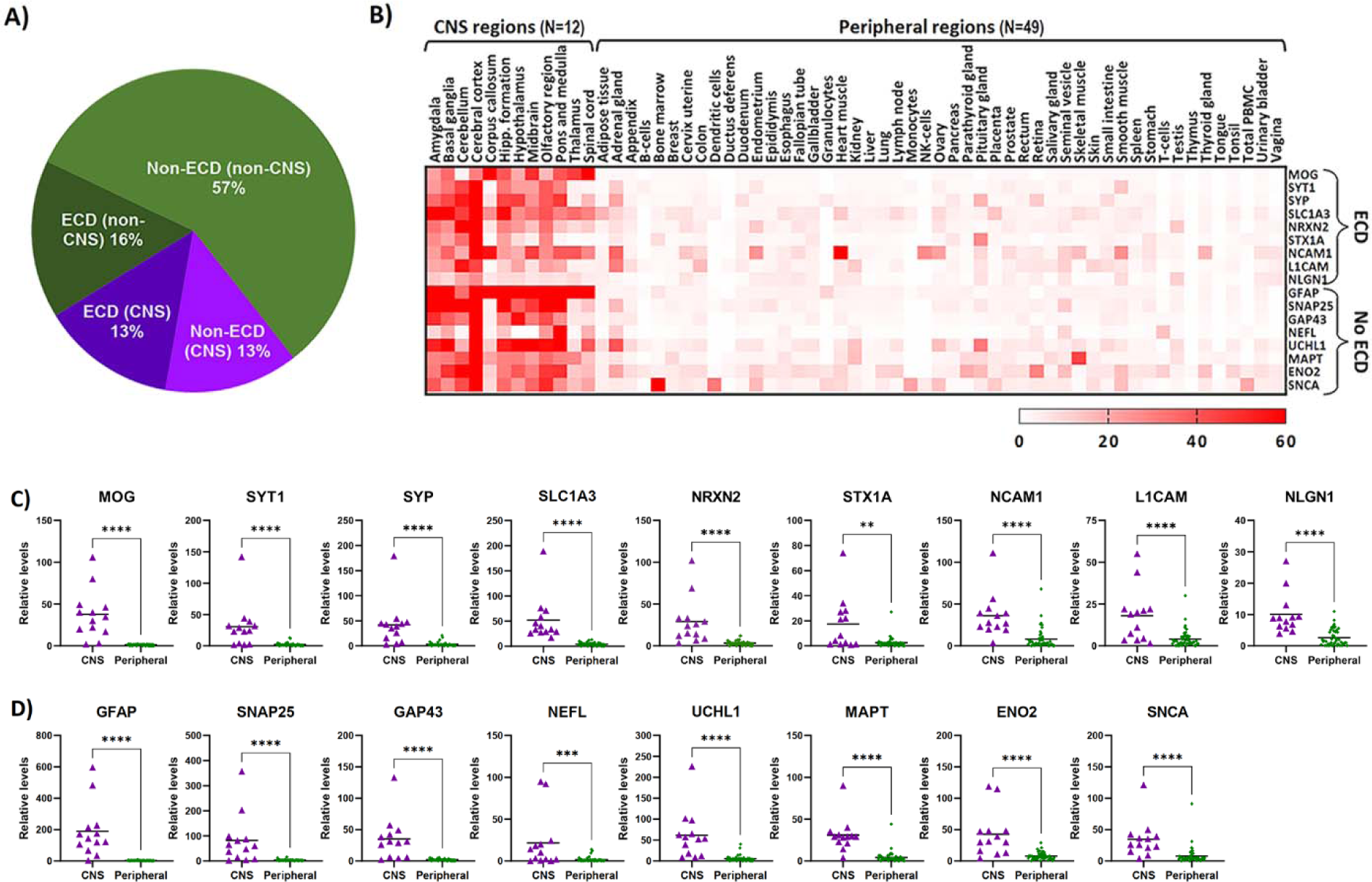
Characterization of proteins in our list **A)** Pie chart showing the distribution of CNS-specific and ECD-containing proteins in our list.Relative gene expression levels across brain (N=12) and peripheral (N=49) regions of 17 proteins identified as CNS-specific and known to be present in EVs presented as both **B)** a heatmap and **C**-**D)** scatter plots. Also indicated are their ECD status, with **C)** showing ECD- containing proteins and **D)** showing non-ECD-containing proteins. The heatmap color scale indicates normalized expression values (nTPM) ranging from 0 to 60. Asterisks in the scatter plots represent statistically significant differences of p<0.0001 (****) or p<0.01 (**) from Mann-Whitney U-test analysis.

**Table 1:**
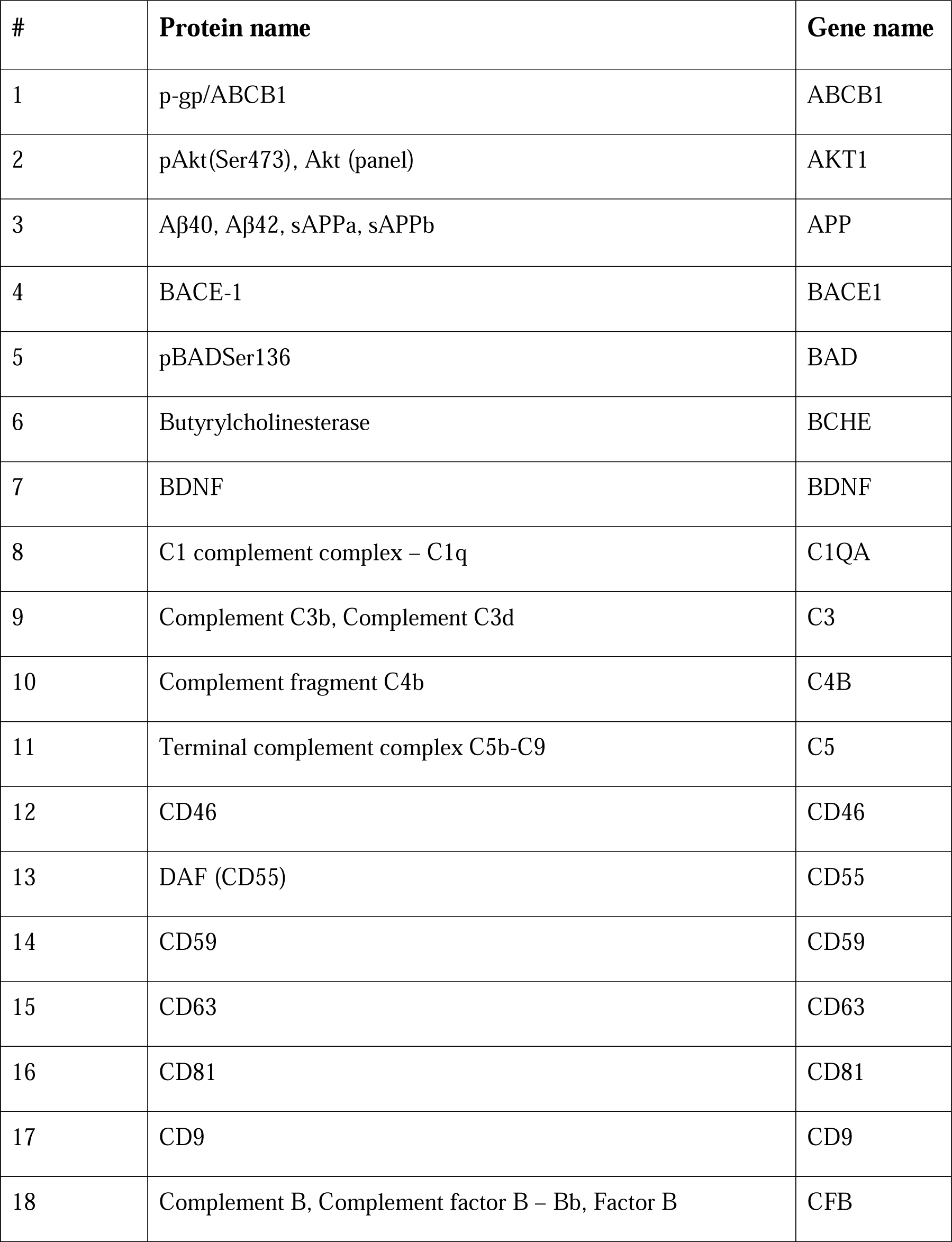
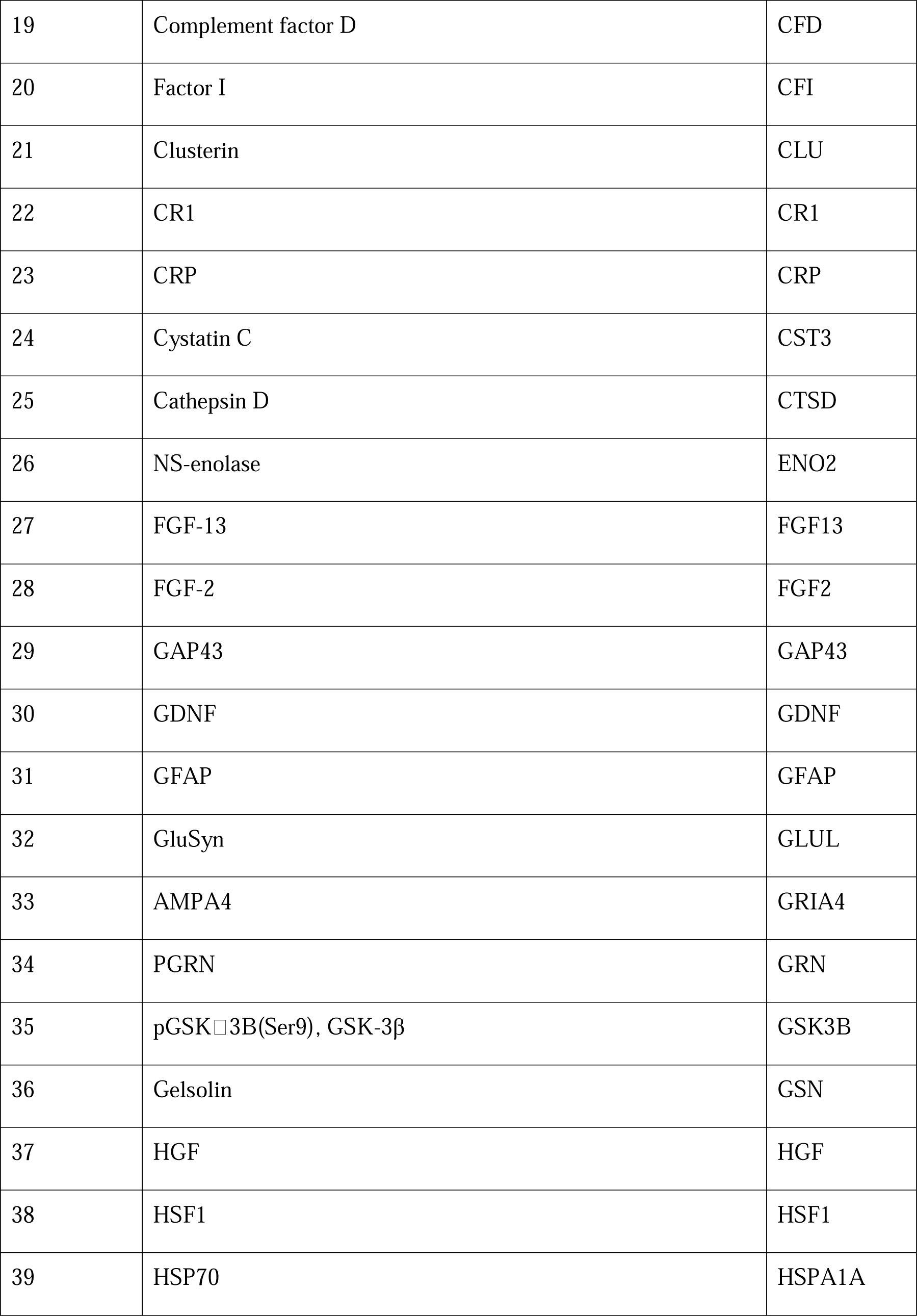
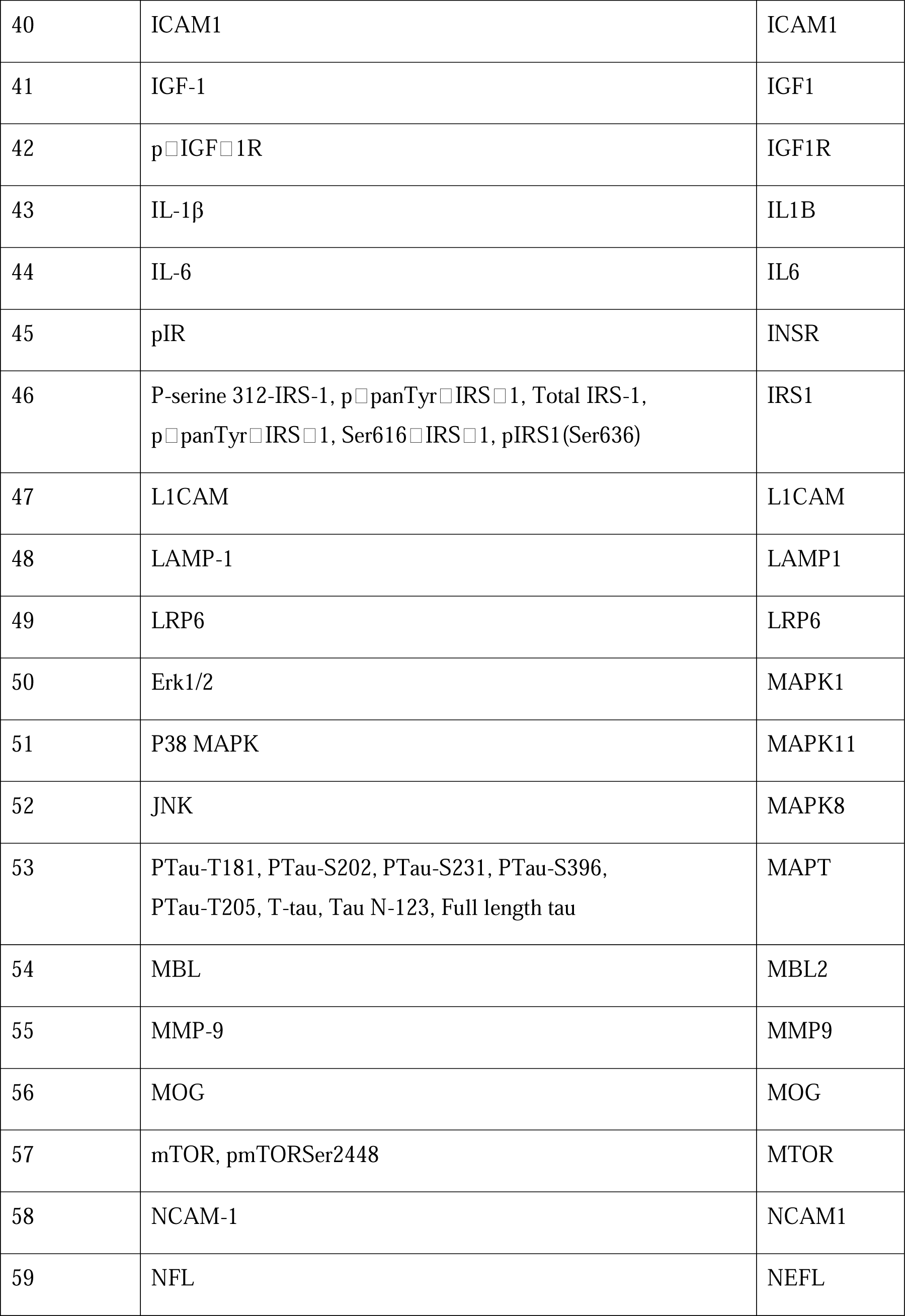
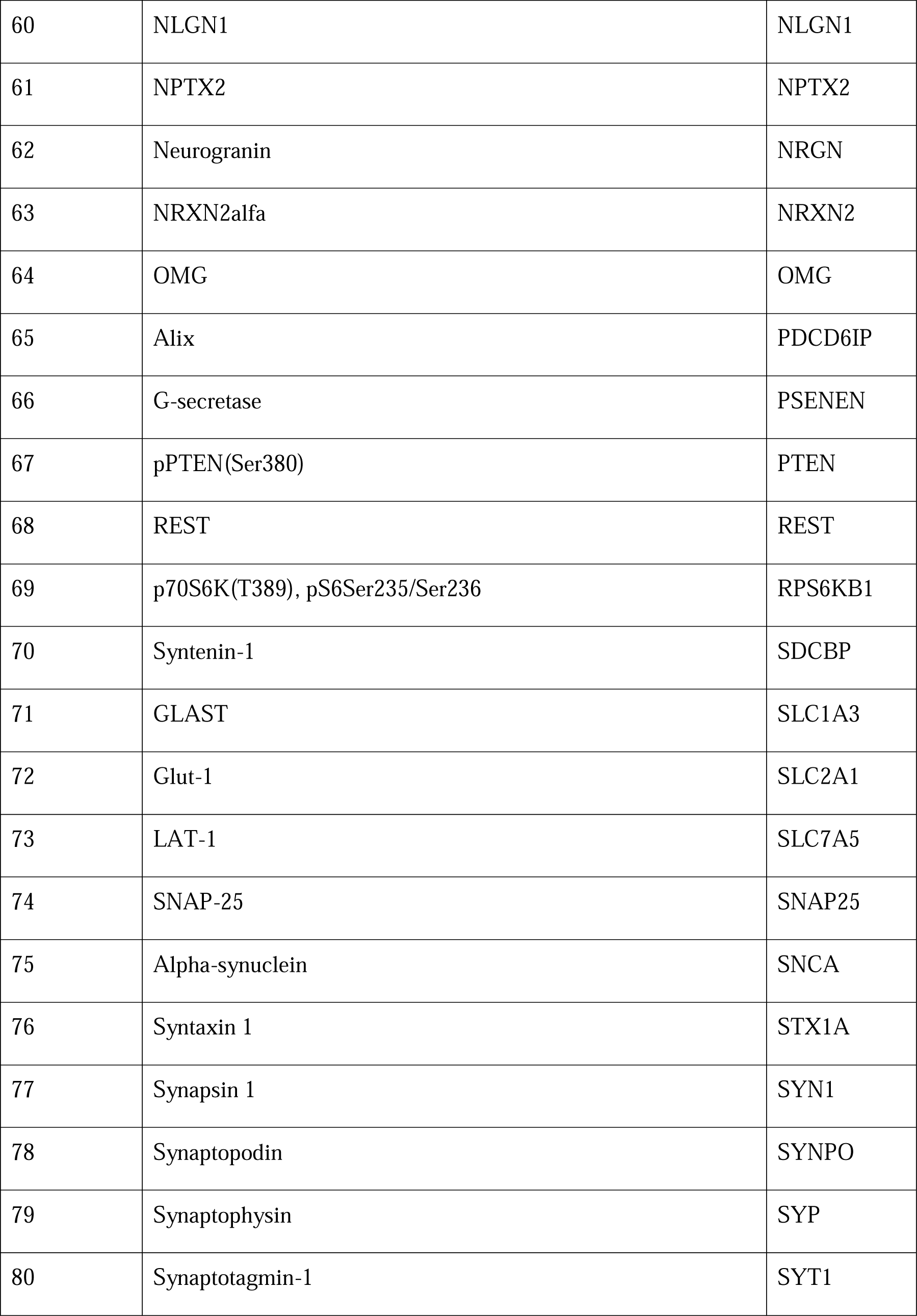
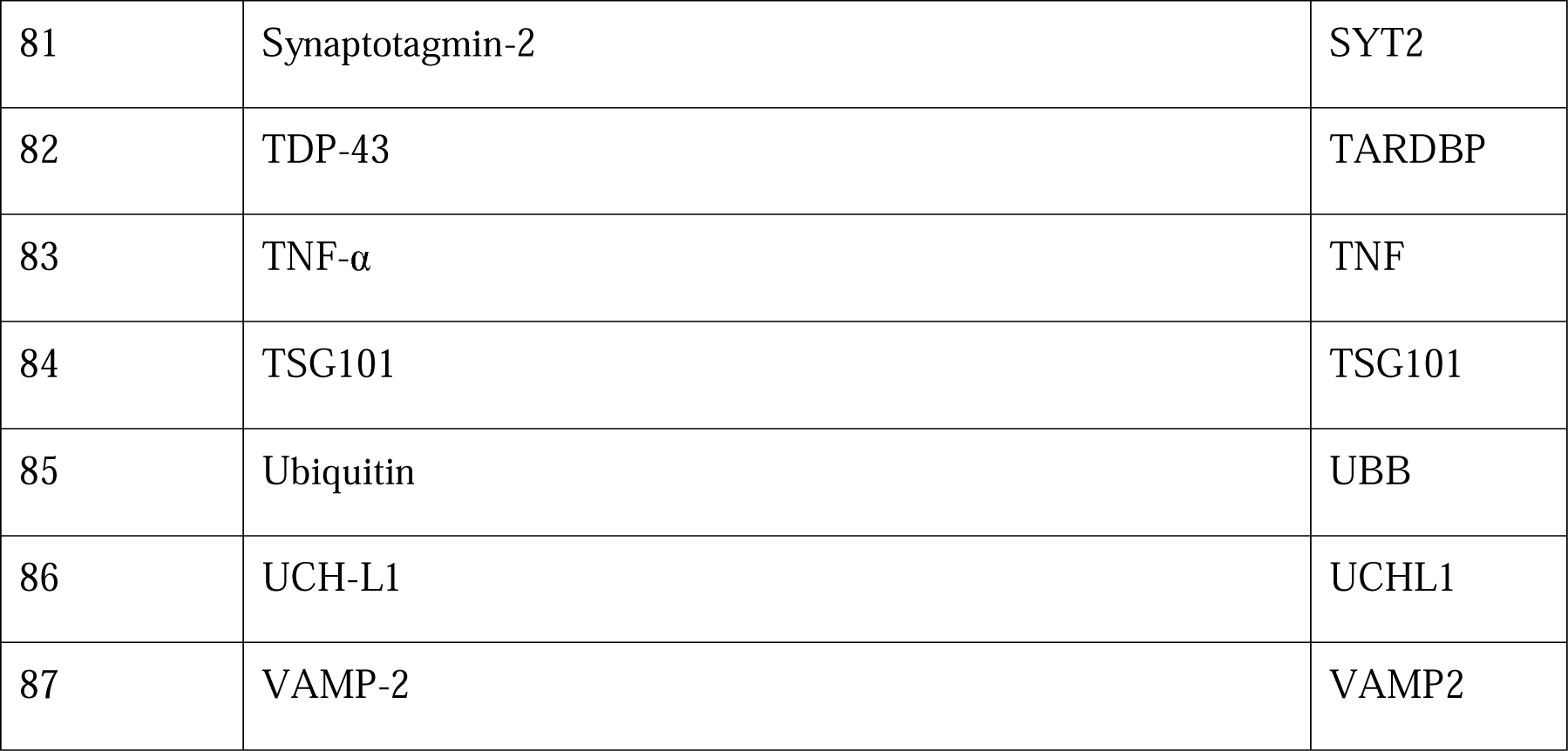
List of 87 unique genes.

Relative expression levels across the CNS and peripheral regions of all 87 genes are shown in Supplementary_Fig_S3.

#### 3.4.1 Brain cell-type specificity of proteins in our list identified as CNS-specific

Of the 22 CNS-specific genes identified, the majority (N=15, 68.2%) were found to be neuron-specific, with the remaining being oligodendrocyte- or astrocyte-specific, or were present in multiple cell types (Fig. 4).

**Figure 4:**
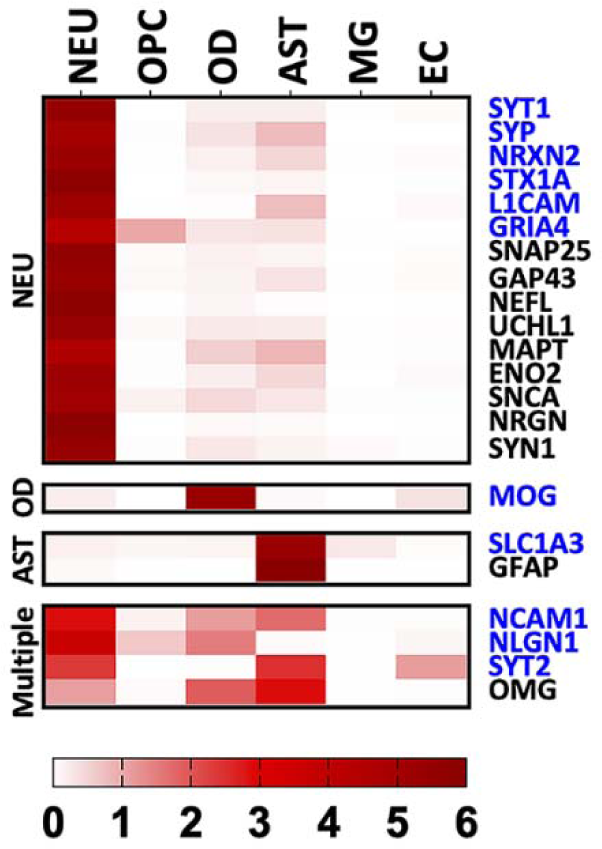
Brain cell-type heatmap The heatmap shows the relative gene expression from a single-cell RNAseq dataset (accession GSE67835, [32]) for each identified CNS-specific protein. Expression in 6 cell-types is shown: neurons (NEU), oligodendrocyte progenitor cells (OPC), oligodendrocytes (OD), astrocytes (AST), microglia (MG), and endothelial cells (EC). The heatmap color scale indicates normalized expression values (relative to average of the 6 cell types) ranging from 0 to 6. Proteins showing normalized values ≥4 were tagged as specific to that cell-type, else they were tagged as present in multiple cell types. Protein names in blue indicate extracellular domain- containing proteins.

## 4. DISCUSSION

Given the increasing usage of BEVs enriched from blood in NDD research, a main goal of the ISTAART-BBB-PIA-EWG was to investigate BEV enrichment methods from plasma and/or serum - a topic often heavily debated by the scientific community. Below, we summarize (a) commonly used experimental choices made across NDD studies for EV and BEV enrichment, and (b) offer ISTAART-BBB-PIA-EWG’s recommendations for CNS-specific alternatives to L1CAM for IP-based BEV enrichment from blood.

### 4.1 Commonly Used Experimental Choices Across NDD Studies

#### 4.1.1 Choice of plasma over serum as starting material

We found that the majority (66.7%) of studies used plasma rather than serum as a starting material to precipitate EVs. This may be due to some of the obvious advantages of plasma, such as (i) the larger volume obtained from a fixed volume of blood, and (ii) no clotting time delay [34], given that rapid processing of blood post-collection is deemed crucial to avoid increasing EV instability with longer (>30 mins) incubation periods [35,36]. Plasma is also considered the most physiological milieu to study blood EVs, taking into account that the number of EVs is higher in serum due to clot-induced platelet vesiculation [35]. Recently, comparisons of fresh versus frozen plasma found no significant difference in protein content of enriched EVs, which may further boost the use of biobanked plasma for EV research [37].

Choice of starting material (i.e. plasma or serum) may, however, impact markers selected for EV validation [38]. While we found tetraspanin CD81 to be the most reported (N=24 studies) EV marker, Karimi et al. [38] recently demonstrated that CD81-positive EVs constituted the rarest subpopulation in plasma and serum. Instead, CD9-positive EVs comprised the majority, with considerable enrichment for CD9- and CD63-positive EVs observed in plasma and serum, respectively [38].

#### 4.1.2 Choice of thromboplastin-D over thrombin for defibrination

We found that thromboplastin-D, and not thrombin, was used by the majority (62.1%) of plasma studies for defibrination. The protease thromboplastin-D converts prothrombin to thrombin during the clotting of blood, while the enzyme thrombin facilitates blood clotting by converting fibrinogen to fibrin. Relative to untreated plasma, pre-treatment with thromboplastin-D was reported to (i) remove clouding factors and prevent aggregation of ExoQuick enriched EVs [39], and (ii) not introduce contaminants, when using human recombinant thromboplastin and enriching for EVs using ultracentrifugation [37], though use of rabbit thromboplastin introduced exogenous tau contaminants [25]. Thrombin pre-treatment, unlike that observed with rabbit thromboplastin, lacked exogenous tau contaminants [25]. Pre-treatment with thrombin, however, significantly lowered Exoquick EV yield compared to untreated plasma [40], with the authors hypothesizing that the induced clotting entrapped a significant number of EVs, thereby leading to an underestimation [40].

#### 4.1.3 Choice of ExoQuick over other approaches for EV enrichment

We found that polymer-based EV precipitation/enrichment on cleared plasma or serum using ExoQuick was used by most (N=32, 82.1%) studies. The observation that 68.8% of the ExoQuick studies followed manufacturer recommendations for incubation time and temperature (60 min, 4°C), as well as centrifugation speed and temperature (1500 x g, 4°C) for precipitation, points to a majority consensus for these parameters. However, (a) the majority of studies (N=19, 59.4%) doubled the manufacturer’s recommendation for ExoQuick for a given volume of plasma or serum, and (b) centrifugation time for precipitation lacked consensus. The impact of these protocol changes to ExoQuick-based EV enrichment requires further investigation.

On the choice of ExoQuick itself for precipitation of EVs from blood, Serrano-Pertierra and colleagues [41] reported that enrichment of plasma-derived EV using ExoQuick was more efficient compared to ultracentrifugation and the Invitrogen kit. Moreover, a recent study, assessing EVs enriched using five commonly used methods, namely, precipitation (ExoQuick ULTRA), membrane affinity (exoEasy Maxi Kit), size-exclusion chromatography (qEVoriginal), iodixanol gradient (OptiPrep), and phosphatidylserine affinity (MagCapture), reported that ExoQuick was a better method for plasma than for conditioned cell media [42]. The study highlighted the importance of selecting an EV enrichment protocol suited to the sample type (e.g. plasma, serum, cell media) [42]. ExoQuick was also found to enrich the most EV-proteins from low plasma volumes (e.g., 250 μL), an important consideration for biomarker studies and clinical trials that lack access to large biosample volumes. However, earlier versions of the ExoQuick kit precipitated both EVs and non-EV-particles (mostly lipoproteins) [42,43], a drawback that may be currently mitigated by the ExoQuick-LP kit, which contains a lipoprotein pre-clearing reagent [42]. It should also be noted that the addition of the immunocapture step with a cell-specific antibody further helps with clearing lipoprotein contaminants [44]. Overall, ExoQuick is considered an easy to perform, fast, reproducible, scalable, and relatively low-cost EV enrichment method [42].

#### 4.1.5 Choice of L1CAM for BEV enrichment

L1CAM or CD171 is a transmembrane protein, known to be widely expressed in neurons [45], and currently serves as the gold standard for enriching NEVs from blood. Numerous studies have demonstrated that NEVs contain Aβ and tau species, relevant to neurodegeneration [23,27,46,47]. Additional work has shown that NEVs and AEVs contain complement proteins, which can accurately predict conversion of MCI to AD [48–50]. Overall, employing cargo analysis of L1CAM-positive BEVs has expanded the field of plasma biomarkers in early AD diagnosis exponentially. However, the specificity of L1CAM itself constitutes the main source of controversy. In 2021, Norman and colleagues [51] reported that L1CAM was not associated with plasma and/or CSF enriched EVs, thereby suggesting that prior BEV biomarker studies were measuring soluble cleaved versions of L1CAM that were not associated with the CNS. Finally, the authors advocated against the use of L1CAM as a marker for enrichment of NEVs [51]. In contrast to the studies investigated in our current work, Norman et al. [51] had used size exclusion chromatography (SEC) to elute their EVs, with L1CAM signal still observable, albeit at lower signal strengths, in fractions where tetraspanin signals (i.e. protein markers of EVs) were also present (e.g., fractions 9-12). Given that EVs are heterogeneous and can range in different sizes, it is possible that L1CAM may be associated with smaller EVs rather than larger EVs. This was not addressed by the authors as the size profile of the EVs eluted in their fractions were not thoroughly investigated. Moreover, given that both soluble and membrane-bound L1CAM can exist together in the same fraction, additional studies robustly differentiating between the two forms are needed. Nonetheless, these findings have sparked a much-needed discussion about the validation and harmonization of protocols in the EV field and require further investigation to counter the numerous studies in support of L1CAM. We found that L1CAM continues to serve as the most used CNS-specific marker for NEV enrichment from blood.

### 4.2 Recommendations for CNS-Specific Alternatives to L1CAM

The issue of BEV specificity and purity arises due to blood containing an admixture of EVs from multiple tissues/cell types. In the current study, we used publicly available RNAseq and EV- databases, to categorize proteins in our list as CNS- and brain-cell-type-specific. We found high expression in 12 CNS compared to 49 peripheral regions for (a) L1CAM, and (b) proteins used for IP-based AEV enrichment, such as GLAST (or SLC1A3) and GFAP. In contrast, CD81 and CD63, well characterized EV markers, demonstrated a wide distribution across both CNS and peripheral regions. Interrogation of the CNS-specific proteins in our list for ECD presence allowed for the identification of potential alternate markers for IP-based BEV enrichment. Specifically, we put forth SYT1, SYP, NRXN2, GRIA4, as potential novel neuronal markers for BEV enrichment using IP-based capture methods. Our findings also suggest that while the proteins NCAM1 and NLGN1 demonstrate a high CNS-specificity, they may not be associated with a specific brain-cell type. Our analyses further revealed candidate proteins with a CNS origin, yet lacking an ECD. These are likely EV cargo proteins that could be used during secondary validation methods. Collectively, the candidate proteins identified here have a wide range of biological functions that may be relevant to neurodegeneration, but require further investigation. Our future work includes validating candidate markers using ultra-sensitive bioassays and proteomic tools for CNS-specific confirmation and novel EV cargo identification.

### 4.3 CONCLUSIONS

Harmonization of protocols are essential steps for blood-enriched BEV biomarker work in AD and other NDDs, and is advocated by the ISTAART-BBB-PIA-EWG. For IP-based BEV enrichment protocols, we have identified steps with obvious variability across studies that require harmonization. We also noted that most studies use L1CAM for enrichment of NEVs. Therefore, additional CNS-specific ECD-containing proteins identified in our study can potentially be used for both IP-based BEV enrichment and as BEV biomarkers. Similarly, CNS-specific cargo proteins can potentially be used for BEV biomarkers.

## ACKNOWLEDGEMENTS

This manuscript was facilitated by the Alzheimer’s Association International Society to Advance Alzheimer’s Research and Treatment (ISTAART), through the Exosome Working Group of the Biofluid Based Biomarkers Professional Interest Area (ISTAART-BBB-PIA-EWG). The views and opinions expressed in this publication represent those of the authors and do not necessarily reflect those of the BBB-PIA membership, ISTAART, or the Alzheimer’s Association. The authors thank the ISTAART-BBB-PIA for their help in initiating this work.

## CONSENT STATEMENT

We confirm that consent was not necessary for this work.

## AUTHOR CONTRIBUTIONS

AB led the study group. AB, ASH, and CNW designed the study. Systematic assessment of protocols were performed by AB, YH, NVT, MFI, AL and CNW. CNS-specificity assessments were performed by ASH and AB. All authors wrote, edited and approved the final manuscript. All authors meet the ICMJE criteria for authorship.

## AVAILABILITY OF DATA AND MATERIALS

The datasets used and/or analyzed during the current study are available from the corresponding author on reasonable request.

## APPENDIX

**COLLABORATORS** - BBB PIA Exosome Working Group (EWG), Established October 2021

1. Alberto Lleo.
2. Amanpreet Badhwar,
3. Charisse Winston,
4. Chi Udeh-Momoh,
5. Henrik Zetterberg,
6. Luc Buee,
7. Maria Florencia Iulita,
8. Michelle M. Mielke,
9. Morvane Colin,
10. Paul Slowey,
11. Rawan Tarawneh,
12. Robert Rissman,
13. Samuel Gandy,
14. Sylvain Lehmann,
15. Tsuneya Ikezu,
16. Aurelie Ledreux,
17. Gagan Deep,
18. Yael Hirschberg

