## Supplemental Figures for "Assessment of brain-derived extracellular vesicle enrichment for blood biomarker analysis in age-related neurodegenerative diseases: An international overview"

### SUPPLEMENTARY MATERIAL

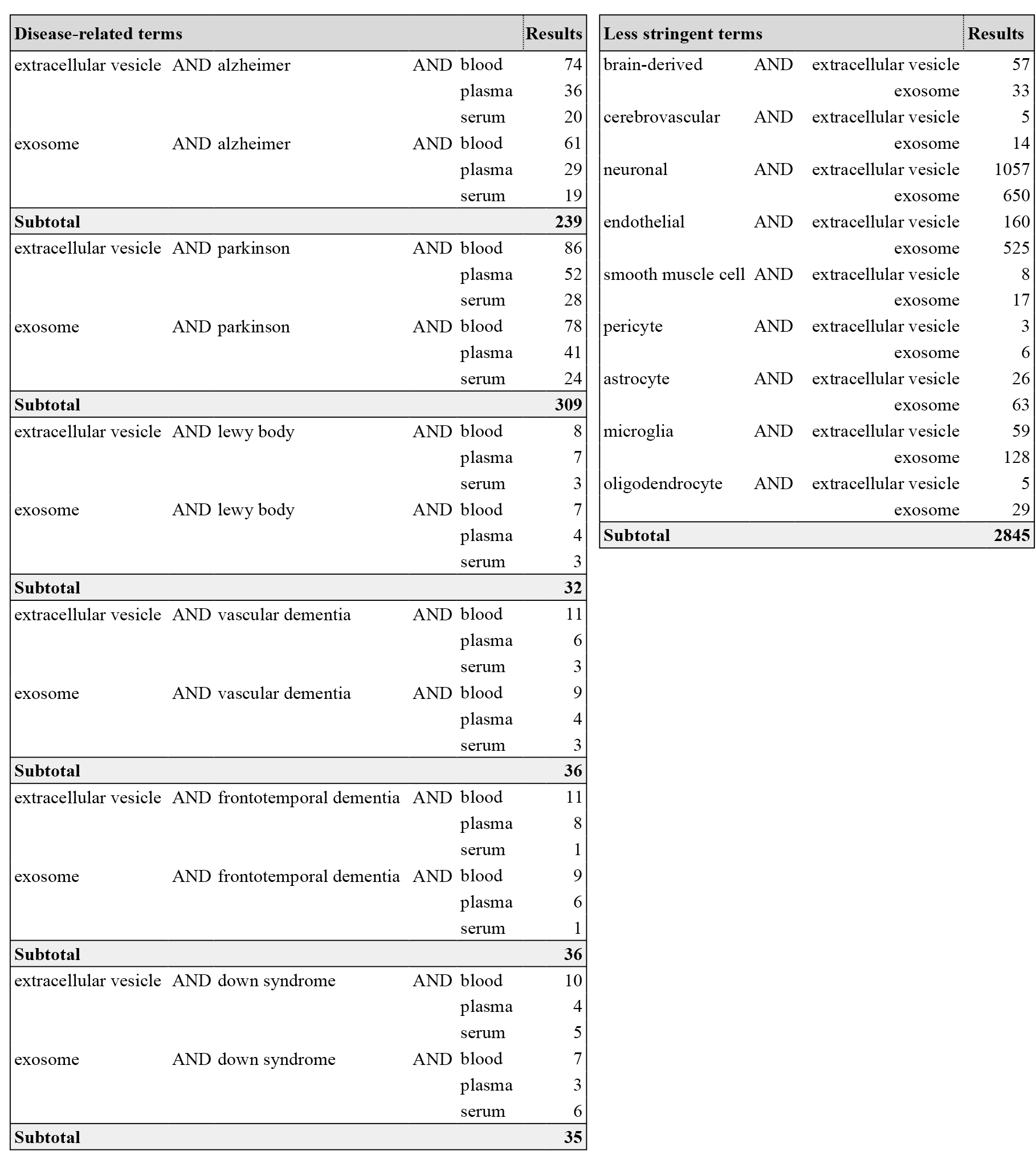
 **Supplementary_Figure_S1**: Search term combinations used.

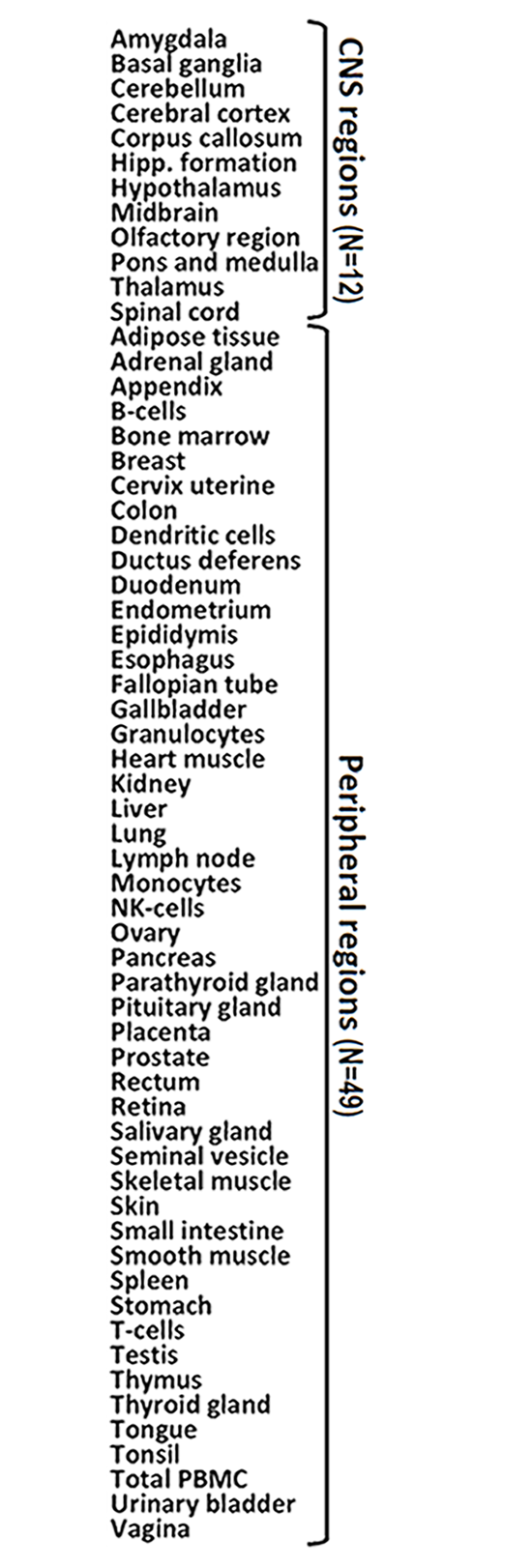

**Supplementary_Figure_S2**: The 61 regions (12 CNS and 49 peripheral regions) with gene expression data.

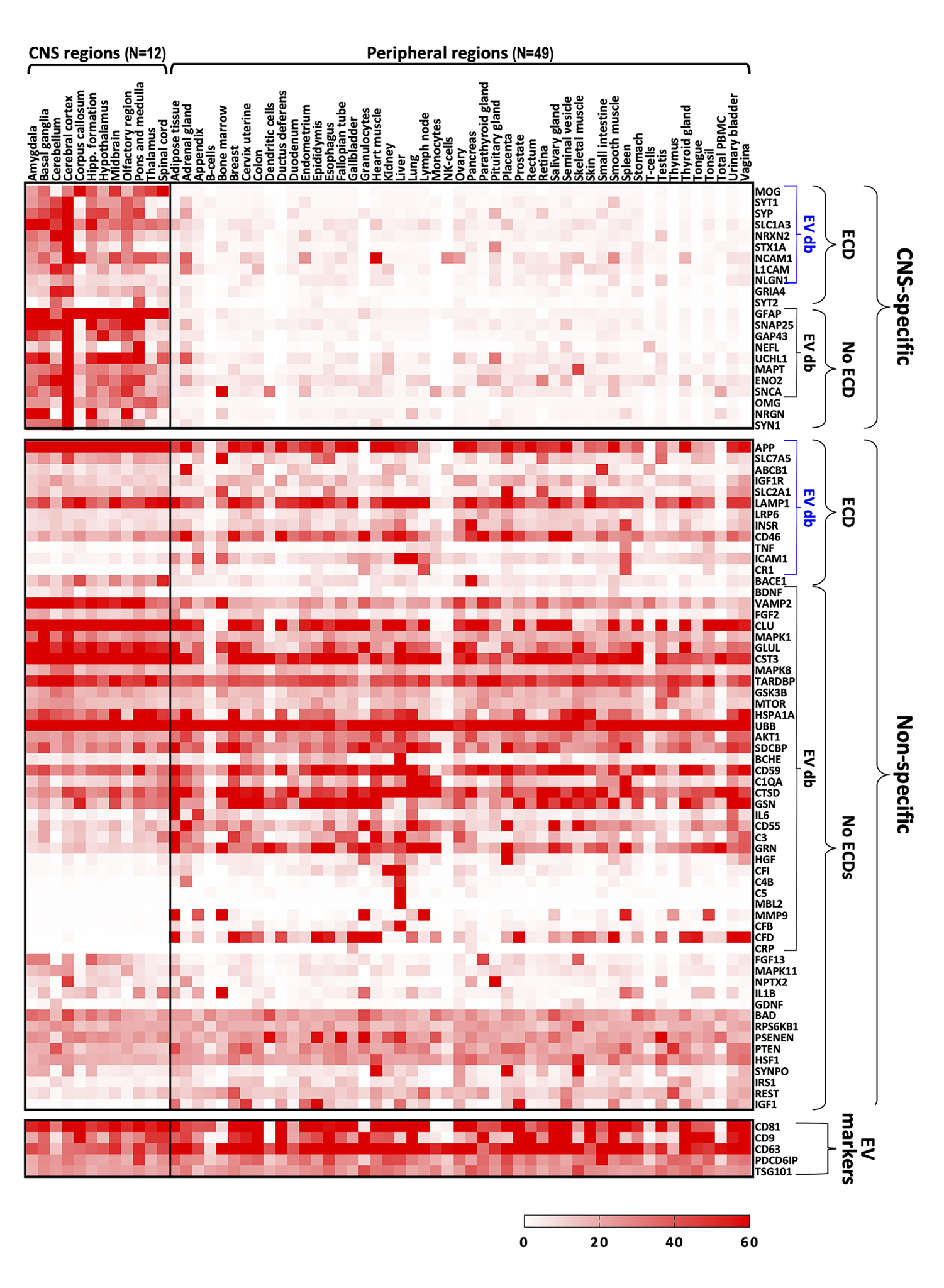

**Supplementary_Figure_S3**: Relative expression levels across the CNS and peripheral regions of all 87 genes shown as a heatmap.
